# Understanding the Dynamics of Fluid-Structure Interaction with an Air Deflected Microfluidic Chip (ADMC)

**DOI:** 10.1101/2022.06.14.496157

**Authors:** Chad ten Pas, Ke Du, Long Pan, Ruo-Qian Wang, Shiyou Xu

## Abstract

A deformable microfluidic system and a fluidic dynamic model have been successfully coupled to understand the dynamic fluid-structure interaction in transient flow, designed to understand the dentine hypersensitivity caused by hydrodynamic theory. The Polydimethylsiloxane thin sidewalls of the microfluidic chip are deformed with air pressure ranging from 50 to 500 mbar to move the liquid meniscus in the central liquid channel. The displacement is recorded and compared with our new theoretical model derived from the unsteady Bernoulli equation. We show that our theoretical model can well predict the ending point of the liquid displacement as well as the dynamics process, regardless of the wall thickness. Moreover, an overshooting and oscillation phenomenon is observed by reducing the friction factor by a few orders which could be the key to explain the dentine hypersensitivity caused by the liquid movement in the dentine tubules.

## A. Introduction

Deformable microfluidics is a unique type of microsystems that possesses at least one deformable sidewall and can be actuated with external applied pressure^1^. This emerging technology has been utilized in automated liquid transport^2,3^, particle/cell sorting^4,5^, and cell mechanics characterization^6,7^. In this project, we demonstrate that this technology can be used to understand the mechanosensitive ion channels in dentine tubules, which cause dentin hypersensitivity problems for over 3 million people each year in the United States^8^.

Dentinal microtubules radiated from pulp wall to exterior dentine-enamel junction (DEJ) contain many microtubules^9^. Most of the dentinal microtubules are filled with non-myelinated terminal fibrils, odontoblastic processes (extension of odontoblast), and dentinal fluid^10^. It is critical to understand how the tooth thermal pain is generated and transmitted to the tooth innervation system through this microtubule structure for oral care industry to design effective tooth pain therapy. The most popular theory of the pain generation and transmission is the hydrodynamic theory^11^, which attributes dental pain sensation to the stimulation of mechano-sensitive nociceptors as a consequence of dentinal fluid movement within dentinal microtubules. Specifically, the thermal deformation of the microtubules would cause a microflow inside the tubules to form a shearing stress to stimulate the odontoblast at the end of the channels. Despite limited computational simulation studies^12,13^, the complicated fluid-structure dynamics has not been validated with fluid dynamics experiments and a theoretical model is needed to understand the fundamental mechanisms. In this project, we leverage the deformable microfluidics to reproduce the microtubules’ flow.

Polydimethylsiloxane (PDMS) is one of the most popular building materials for deformable microfluidics with the advantages such as high deformability, biocompatibility, and stability^14^. We recently showed that the mechanical properties of the PDMS-based deformable microfluidics enable controlled microparticle capture and release^15^. However, the past studies focused on the fluid-structure interaction in steady flows^16–20^. In comparison, systematical studies of the fluid-structure interaction in transient flows are much less. For example, Whittaker et al. developed a theoretical model of tube oscillation with elastic wall^21^. Nevertheless, a dynamic theoretical model that can be validated against transient experimental observation has not been fully established.

In this work, we construct an Air Deflected Microfluidic Chip (ADMC) with PDMS and study the fluid displacement by inputting the air pressure on the deformable sidewalls. The meniscus height change is characterized by a commercial available goniometer and a custom designed LabVIEW program is used to process the collected images. The present study is designed to fill the knowledge gap in fluid-structure interaction in transient flows by performing microfluidics experiment and develop a theoretical model that can match and explain the fluid dynamics observation, thus establishing a key step for the understanding of dentine hypersensitivity in microscale.

## B. Experiments

### Negative mold fabrication

The negative mold was fabricated using an ELEGOO Saturn mono-stereolithography (MSLA) 3-D printer with Accura 25 resin (**Figure 1a-i**). Due to the high hydrophobicity, the Accura 25 resin allows for demolding of cured PDMS without salinization. High aspect features such as large length versus small width/height and large height versus void geometry were achieved at a resolution of 50 μm. The complete mold was constructed out of 3-D printed sidewalls with grooves to hold 4 typical glass microscope slides. The wall structure can be freely placed on the negative mold with a loose tolerance. Use of a sealing membrane such as aluminum foil secured by tape avoids PDMS leaking during mold casting and curing. The mold and wall structure are reusable provided cleaning between each casting. In this work, we tested the samples with an aspect ratio (height/thickness) ranging from 1.5:1, 2:1, and 3:1.

**Figure 1.**
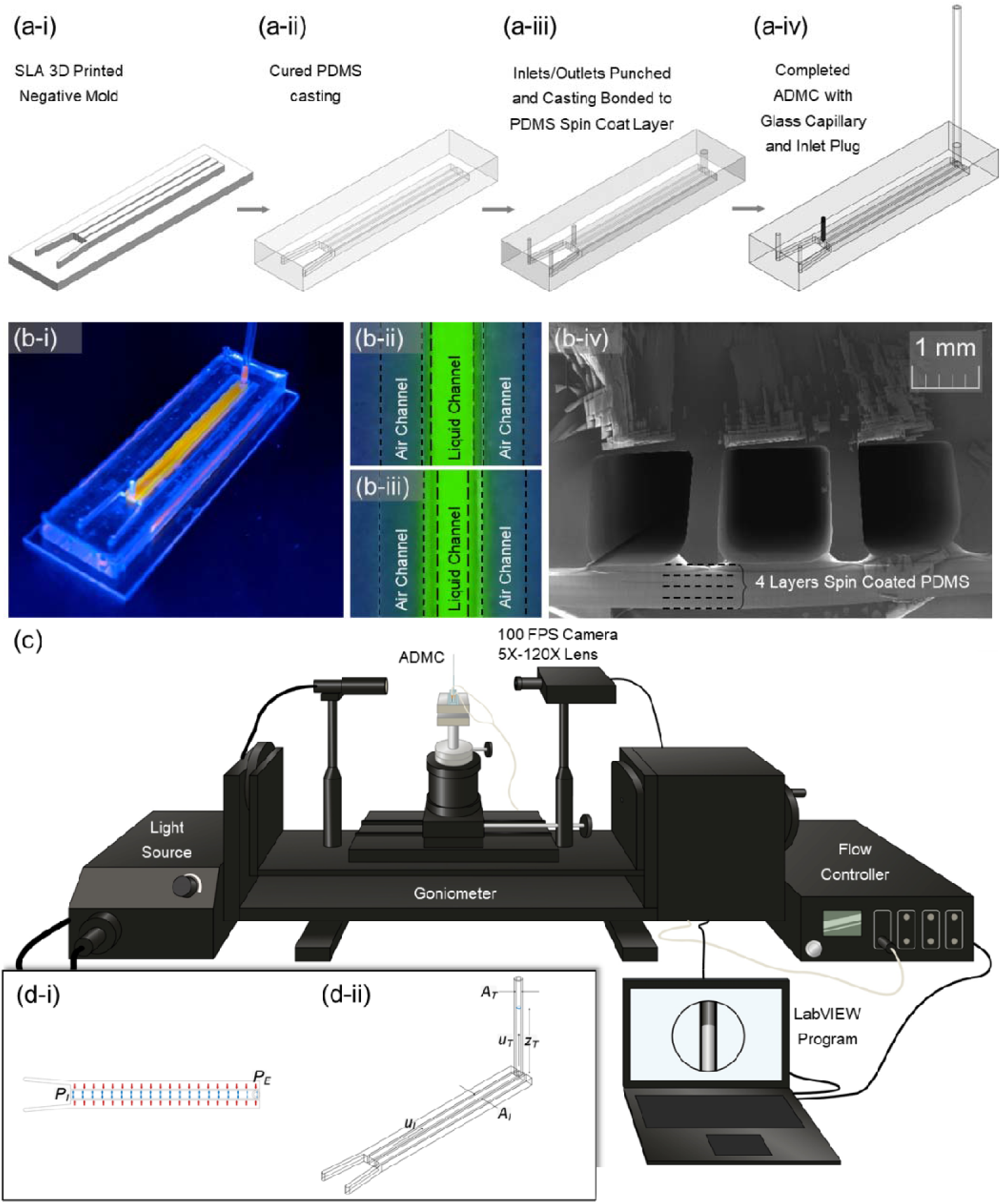
(a) The fabrication process of the ADMC: (a-i) SLA 3-D printed negative mold. (a-ii) PDMS casting on the SLA mold. (a-iii) Inlets and outlet of the channels are punched. The chip is bonded with PDMS to a 1 mm thick spin coated PDMS layer. (a-iv) The central channel has a glass capillary inserted as a sight glass. After filling the central channel with distilled water, the inlet is plugged. (b-i) Photograph of the ADMC with orange fluorescent dye to distinguish between the liquid and air channels. (b-ii) and (b-iii) Before and after application of pressure to outside channels with green fluorescent dye. The thin wall between deflects and displaces fluid. (b-iv) Cross-section of the ADMC with a 500 μm sidewall. (c) The experimental setup – goniometer fitted with a 5X-120X magnification c-mount lens. The pressure is supplied using an ElvFlow controller. The camera and the flow controller are controlled using a LABVIEW program developed to collect images in synchrony with pressure sensor readings. (d) Schematics of the theoretical model.

### Microfluidic chip fabrication

The standard ratio of PDMS base to crosslinker of 10:1 was used in accordance with manufacturer recommendations. We casted 10 mL of PDMS into the negative mold and a standard 25 psi vacuum desiccator was used to remove air bubbles from the uncured PDMS, paying special attention to removing air bubbles in void gaps (**Figure 1a-ii**). The casted PDMS and mold were placed in a free convection oven at 45°C for at least 4 hrs. Low temperature is required to avoid entering glass transition and deformation of the negative mold. After demolding, a biopsy punch (diameter: 1.5 mm) was used to form inlets and outlets in the desired channels (**Figure 1a-iii**). For chip completion, we used a spin coated PDMS layer of approximately 1 mm in thickness as a bottom layer of the chip. To achieve this 1 mm layer, we spin coated four layers of PDMS at 350 RPM for 1 min per layer, curing in between each layer to achieve a layer of PDMS with an approximate thickness of 1 mm. Then, a very thin film of PDMS was spin coated on the cured 1 mm layer at 1000 RPM for 1 min as an adhesive layer between the spin coated PDMS and the casted PDMS chip. Finally, a glass capillery and an inlet plug were inserted into the ADMC chip (Figure **1a-iv**). A photograph of the dye filled ADMC sample is shown in **Figure 1b-i**. As shown in **Figure 1b-ii** and **1b-iii**, after applying air pressure to the air chambers, the PDMS thin walls are deformed and squeezing the fluorescence dye. The cross-section of the chip is shown in **Figure 1b-iv**, clearly showing the stacked PDMS layers at the bottom.

### Measurements of applied pressure and fluid displacement

The experimental setup is shown in **Figure 1c**. Reservoir pressure was supplied to the system using lab air connected through a Drierite gas purifier (desiccator) inline to an Elve Flow flow controller to keep the air supply dry and clean. The flow controller delivers reservoir pressure with a resolution of 100 μbar to the gas channels on the ADMC through rigid pneumatic tubing. The flow controller was outfitted with pressure transducers to measure applied pressure. The liquid meniscus height in the glass capillary tube was measured using a 5X-120X microscope lens attached to a 100 FPS goniometer camera. The acquisition sampling time was set to 60-75 Hz. A LabVIEW program was designed to synchronize the frame collection with the pressure measurements.

### Detecting fluid displacement by image processing

An interactive routine was created to streamline and control processing of the image frames and pressure data. MATLAB was used to track the position of the liquid meniscus by means of image processing. Each image was converted into an edge image using a Canny edge detection function, where the contrasting edges were converted to white lines on a black background. A sensitivity threshold was set between 0.2 - 0.6 and tuned depending on light intensity. To convert the image to engineering units, we performed a centroid calculation to determine the displacement of the meniscus of that frame relative to the first. A pixel conversion factor was calculated for each dataset based on the known thickness of the glass capillary in observation. Each glass capillary has an outside diameter of 2 mm ± 0.1mm. Compiling each of the frame data points together yielded the fluid dynamic response of our chip.

### Theoretical model

A theoretical model was developed to understand the dynamics of fluid-structure interaction and to verify the experimental results. The model (**Figure 1d-i and 1d-ii**) generally includes two sections – the pressurized channel and the vertical tube at the end of the channel. Applying the unsteady Bernoulli equation to the device,

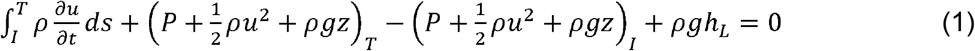

where *P* is the pressure, *ρ* is the fluid density, *μ* is the velocity, *t* is time, *z* is the height of the fluid (*z_I_*=0 and *z_T_* is the height of the free surface in the vertical tube), and *h_L_* is the head loss in the flow, which generally takes account of the major and minor losses in the device. The subscription *T* and *I* denote the variables in the vertical tube and the horizontal pressurized channel. The total fluid in the device is conserved, therefore,

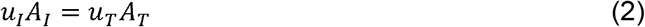

where *A_I_* and *A_T_* are the cross-sectional areas of the pressurized channel and the vertical tube. To describe the tube deformation under the transmural pressure, the tube law is applied^22^, i.e.,

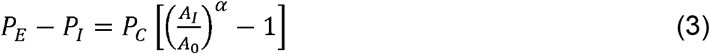

where *A_O_* is the initial cross-sectional area, *P_E_* and *P_I_* are the pressure outside and inside the channel, *P_C_* is a deformation coefficient, and *α* is the exponent depending on the shape and materials of the channel. Various tube laws are proposed in the literature and the equation above is from the most popular one used in flexible tubes. Assuming the velocity in the pressurized channel is uniform and inserting the laminar Darcy-Weisbach equation, i.e.,

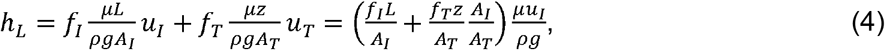

where *L_L_* and *L_A_* are the length scales of the channel length and cross-sectional characteristic length such as the tube diameter, we obtain,

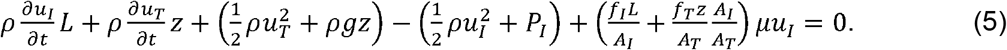

Normalizing the equations yields,

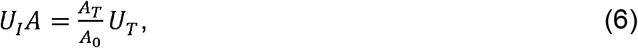

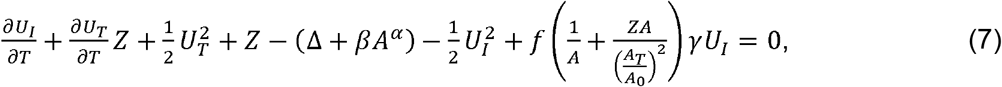

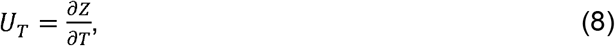

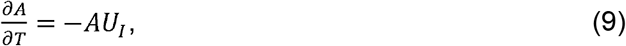

where 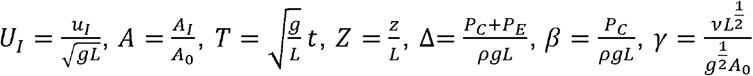, and assume *f* = *f*_I_ = *f*_T_.

This system of equations was solved using the ode15i function in MATLAB. The initial velocity was assumed zero and no deformation was assumed at *t*=0.

## C. Results and Discussions

**Figure 2a** shows the raw images labelled with 1, 2, 3, and 4 correspond images captured at 0 s, 0.10 s, 2.34 s, and 23.77 s, respectively. Height values that were calculated are shown alongside the time values. The frames from **Figure 2a** were processed further using a Canny edge detection algorithm. In addition to the canny edge detection, the images are cropped to omit extra and unneeded data, as shown by the white rectangle in the frames. **Figure 2c** shows the cropped areas from **Figure 2b**. The top and bottom edges of the liquid meniscus are averaged together by centroid calculation to output the liquid surface height relative to the first frame. The average meniscus height is shown in magenta in **Figure 2c**. Each image in **Figure 2a**, **2b**, and **2c** correspond to the data points marked in **Figure 2d**. **Figure 2d** shows a sample data set from a 500 μm wall thickness device with 200 mbar of pressure applied.

**Figure 2.**
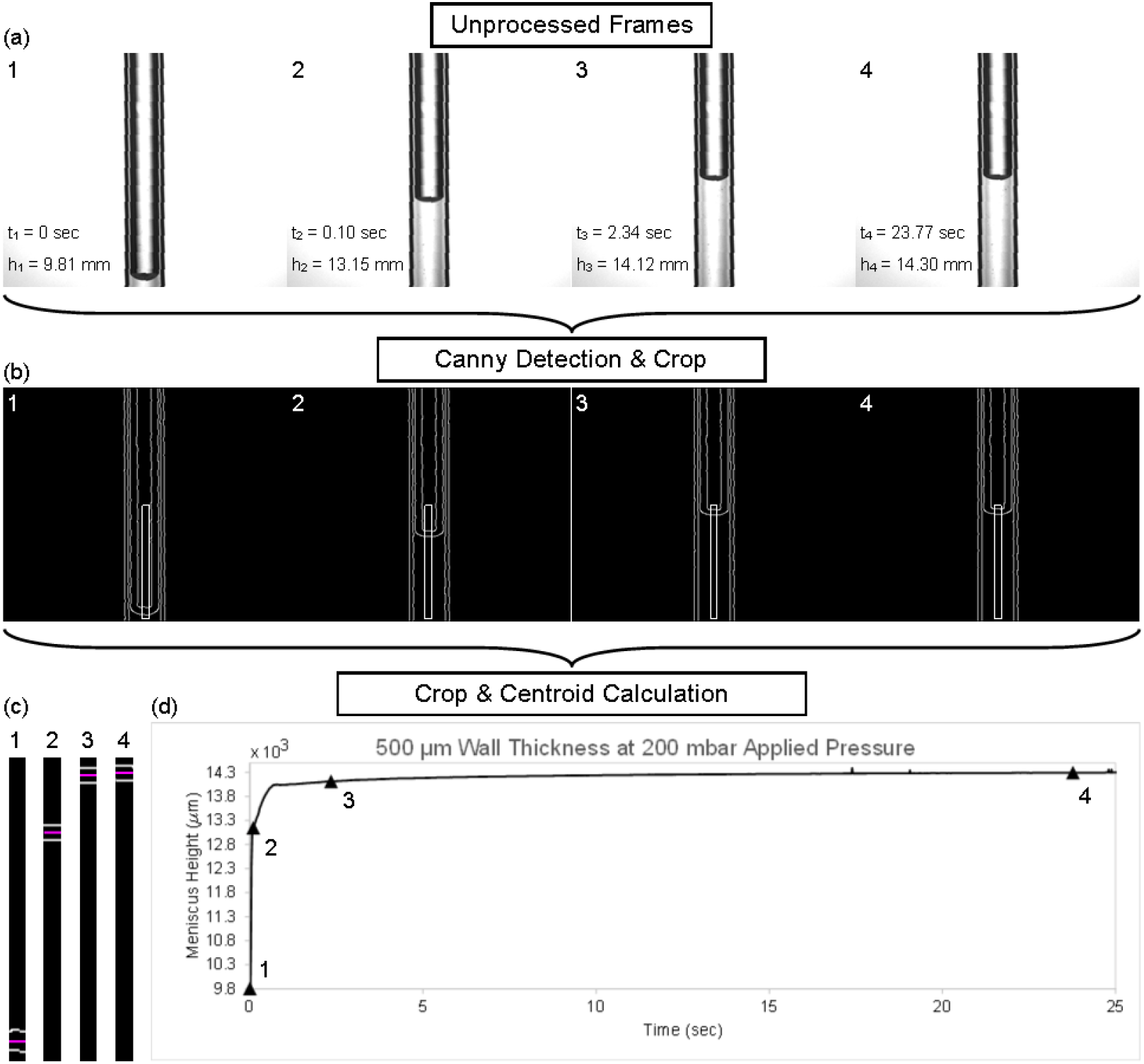
(a) Frames are collected at 60-10 Hz and are rotated so the 1.2 mm ID glass capillary is upright. The frames need to be high contrast for (b) Canny edge detection. The meniscus at initial and maximum height is (c) cropped on each image. Centroid calculations are performed to find the meniscus height at each frame. (d) Each frame height is compiled together to form an output plot.

The measured meniscus height versus time under various air pressure is shown in **Figure 3**. For 500 μm thick sample (**Figure 3a-i** and **3a-ii**), the meniscus shows a sharp increase within the first 10^th^ of s, regardless of the input air pressure. In the second phase, the meniscus gradually rises over the course of ~0.5 s. Then, the meniscus reaches to a steady state without showing much change from 1 to 10 s. With 500 mbar air pressure, the meniscus height reaches to ~12,000 μm in contrast to 1,000 μm for 50 mbar. As shown in **Figure 3b** and **3c**, the 750 μm and 1,000 μm samples follow a similar trend with the 500 μm sample. However, the maximum meniscus height only reaches 6,500 and 1,700 for 750 μm and 1,000 μm samples, respectively.

**Figure 3.**
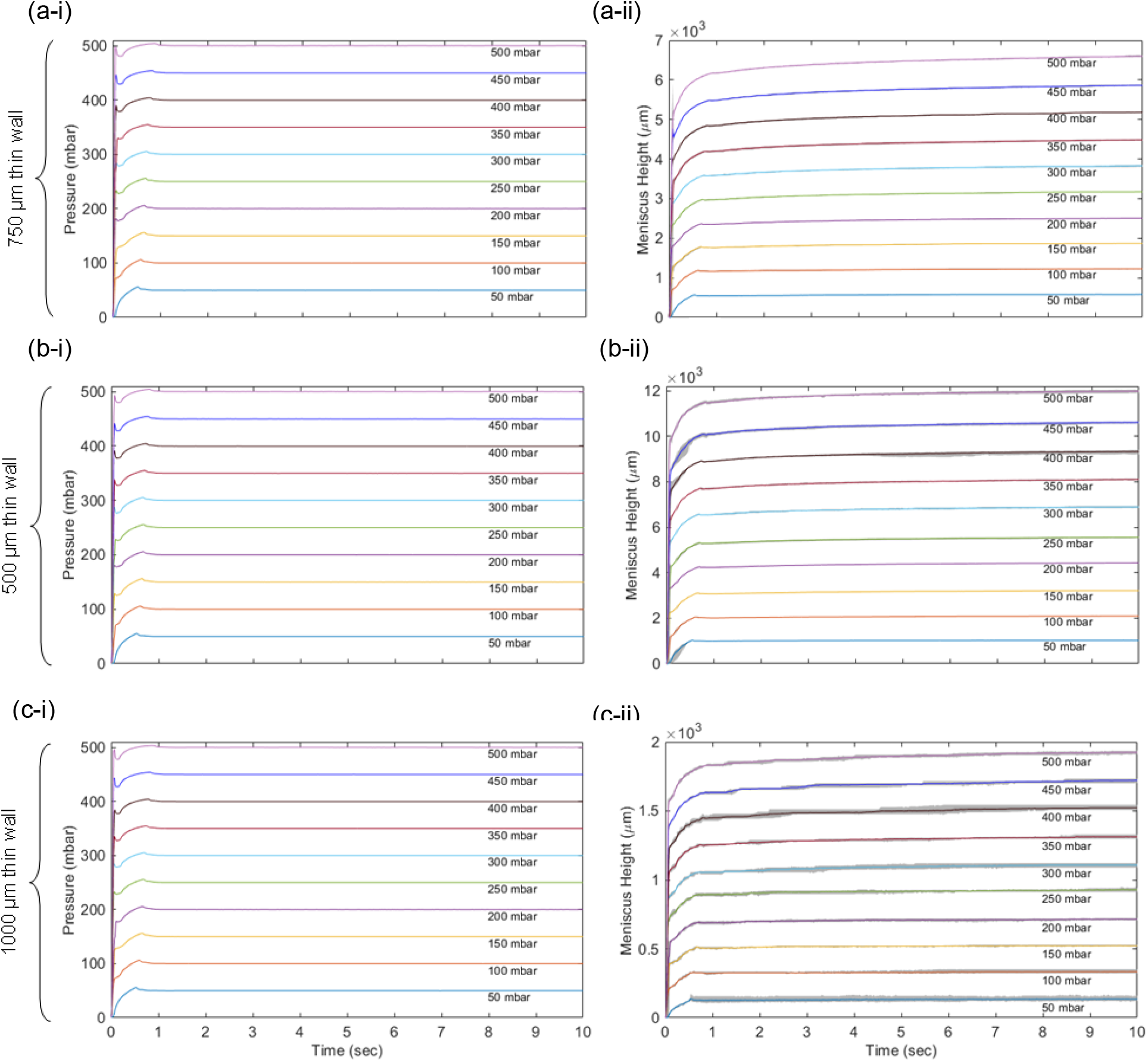
Input pressure and output meniscus height of three different thin wall chip geometries: (a) 500 μm. (b) 750 μm. (c) 1000 μm. The meniscus height response increases linearly with application of pressure and reduces as we increase the thin wall thickness.

The experimental results and simulation comparison for 500, 750, and 1,000 μm samples are shown in **Figure 4a**, **4b**, and **4c**, respectively. Our theoretical model well captures the dynamics of the second and third phases of the dynamic process, regardless of the wall thickness. Specifically, the heights at the end of Phase 1 were used as the starting points of the modeling and the other used parameters in the theoretical model to compare with the experimental results are listed in **Table 1**. The theoretical model does not only compare well with the ending points of the experimental results but also with the dynamics. The change of the rising rate and the magnitudes are well reproduced. This indicates that the theoretical model can, to an extent, reflect the complicated dynamics in the experiments in addition to the steady state results that are dictated by the mass conservation before and after the deformation.

**Figure 4.**
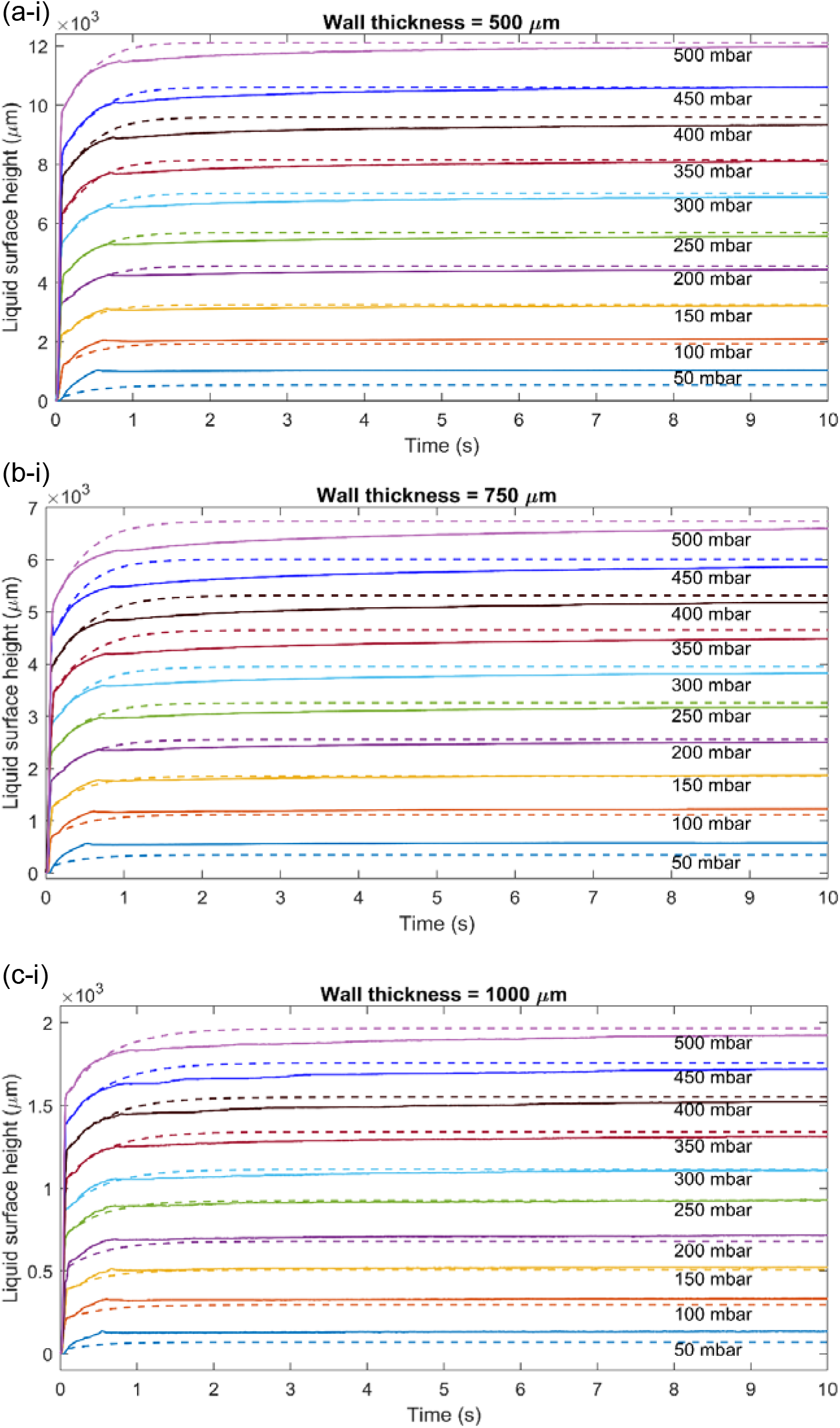
Experimental trials (solid lines) overlayed with simulated (dashed lines) theoretical model for (a) 500 μm. (b) 750 μm. (c) 1,000 μm.

**Table 1.** The parameters used in the theoretical model to compare with the experimental results.

| Wall thickness ($\mu\text{m}$) | $\alpha$ | $P_c$ | $f$ |
| --- | --- | --- | --- |
| 500 | -92 | $2.2 \times 10^4$ | $5 \times 10^5$ |
| 750 | -65 | $6.3 \times 10^4$ | $8 \times 10^5$ |
| 1000 | -21 | $1.06 \times 10^6$ | $5 \times 10^6$ |

Much effort was also made to capture the dynamics of Phase 1, but it turns out that the meniscus height rose too quickly to compare with the theoretical model. This indicates that the process in Phase 1 was governed by a different mechanism beyond the framework of the hydrodynamics described in the theoretical model. We suspect this process was formed due to the pre-tension during the fabrication process – when the pressure was applied, the pre-deformation played a significant role to accelerate the meniscus’ rising. A further investigation with different fabrication stress is needed to resolve this issue.

The last process of the experiment still shows a slow increasing trend, which is not captured by the theoretical model. This difference might be due to the response time of the structure deformation, while in the theoretical model the deformation was assumed to be instantaneous. In addition, the streamwise non-uniformity might be attributed to the difference.

An observation can be made in the parameters used in the theoretical model. The tube law’s *α* and *P_c_* increases with the channel wall thickness. This is expected as the thicker wall leads to less flexible cross-sectional changes. The friction factor’s change is not expected as the wall thickness should not change the roughness of the channel surface. This might be attributed to the minor loss at the connection between the channel and the tube.

As mentioned earlier, the present theoretical model assumes that 1) the channel deforms uniformly; 2) surface tension is neglectable at the meniscus in the tube; 3) the minor loss at the connection between the channel and the tube could be ignored; 4) the tube was assumed vertical; 5) the deformation of the channel was assumed to be instantaneous; and 6) the total fluid mass was assumed conserved. These assumptions may contribute to the overall uncertainty of the theoretical model’s performance and future studies should be targeted to examine these over-simplifications.

Interestingly, if the friction factor is reduced by a few orders to allow stronger inertia in the flow dynamics, we observe an overshooting and oscillation process (**Figure 5**). This is consistent with the 3-D computational fluidic dynamic studies by other groups^12,23^. The oscillation could be the key to the tooth induced by thermal triggers and our theoretical model well predicts its presence under the current framework. A further comparison with the existing numerical model is required to investigate the complicated mechanisms revealed in this study and this could lead to resolve the mystery of the particle-contained tooth paste – particle-containing tooth paste was observed effective in reducing dentine-hypersensitivity^24^. However, this mechanism has not been fully understood. The particles contained in the toothpaste could block the open tubules in damaged teeth to reduce the oscillation magnitude of the flow to reduce the pain. This hypothesis has not been tested in simulation or experiments. Our study provides a theoretical framework and experiment setup to investigate the impact of the boundary condition of the tubule flow on the flow dynamics and thus shed lights on this issue. A further study is planned to move forward in this direction. Another knowledge gap revealed in this study is the scale of the model. As the friction factor could determine the presence of the oscillation of the flow and the scale of multiphase flow determines the physics as shown in other studies^25^, a detailed study to determine the parameters in the lab scale and the microtubule scale is required to fully address the model scaling problem.

**Figure 5.**
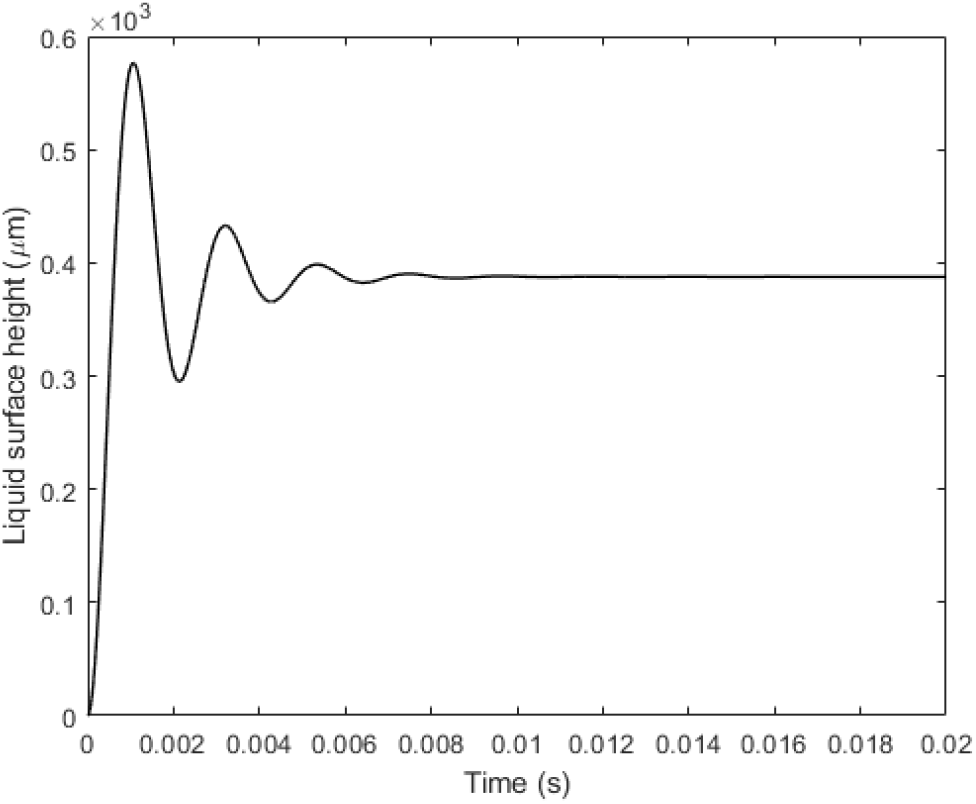
Observed liquid meniscus oscillation at the first 0.008 s in our theoretical model.

